# Progressive Multiple Sequence Alignment with the Poisson Indel Process

**DOI:** 10.1101/123513

**Authors:** Massimo Maiolo, Xiaolei Zhang, Manuel Gil, Maria Anisimova

## Abstract

Sequence alignment lies at the heart of many evolutionary and comparative genomics studies. However, the optimal alignment of multiple sequences is NP-hard, so that exact algorithms become impractical for more than a few sequences. Thus, state of the art alignment methods employ progressive heuristics, breaking the problem into a series of pairwise alignments guided by a phylogenetic tree. Changes between homologous characters are typically modelled by a continuous-time Markov substitution model. In contrast, the dynamics of insertions and deletions (indels) are not modelled explicitly, because the computation of the marginal likelihood under such models has exponential time complexity in the number of taxa. Recently, Bouchard-Côté and Jordan [PNAS (2012) 110(4):1160–1166] have introduced a modification to a classical indel model, describing indel evolution on a phylogenetic tree as a Poisson process. The model termed PIP allows to compute the joint marginal probability of a multiple sequence alignment and a tree in linear time. Here, we present an new dynamic programming algorithm to align two multiple sequence alignments by maximum likelihood in polynomial time under PIP, and apply it a in progressive algorithm. To our knowledge, this is the first progressive alignment method using a rigorous mathematical formulation of an evolutionary indel process and with polynomial time complexity.

## 1 Introduction

Multiple sequence alignments (MSAs) are routinely required in the early stages of comparative and evolutionary genomics studies. Not surprisingly, accuracy of MSA inference affects subsequent analyses that rely on an MSA estimates [13]. MSA estimation is among the oldest bioinformatics problems, yet remains intensely studied due to its complexity (NP-hard [5, 1, 12]). The progressive alignment approach has allowed to reduce the overall computational complexity to polynomial time by breaking the MSA problem into a series of pairwise alignments guided by a tree representing the evolutionary relationship of sequences. Today most popular alignment programs employ the progressive approach (e.g., ClustalW [10], MAFFT [6], MUSCLE [4], PRANK [7] and T-Coffee [3, 9] among others).

All state of the art MSA programs nowadays use an evolutionary model to describe changes between homologous characters, providing a more realistic description of molecular data and thus more accurate inferences. However, a mathematical formulation of the insertion-deletion (indel) process still remains a critical issue. Describing the indel process in probabilistic terms is more challenging: unlike substitutions, indels often involve several sites, vary in length and may overlap obscuring the underlying mechanisms. Instead, the popular PRANK program adopts a pragmatic approach; it uses an outgroup to distinguish insertions from deletions during the progressive alignment procedure, so that they are penalised differently [8]. As a result, PRANK produces exceptionally accurate alignments, notably with densely sampled data and given an accurate guide tree. Still the method lacks a mathematical model describing the evolution of indels. Indeed, the computation of the marginal likelihood under the classical indel model TKF91 [11] is exponential in the number of taxa due to the absence of site independence assumption.

A recent modification of the TKF91 describes the evolution of indels on a phylogenetic tree as a Poisson process, thus dubbed the Poisson indel process or the PIP model [2]. Consequently, standard mathematical results, particularly the Poisson thinning, allow to achieve linear time complexity for computing the joint marginal probability of a tree and an MSA. This includes analytic marginalisation of unobservable homologous paths which occur whenever an ancestral character is inserted and subsequently deleted, and consequently cannot be detected in the extant sequences. For a given MSA and a tree, a likelihood score under PIP can be computed in linear time. This score can be used to find the maximum a posteriori tree-alignment solution. Remarkably, this breakthrough allows for a necessary rigorous way of combining models of substitutions and indels, and a tractable computation of the marginal likelihood function. Nevertheless, at the moment the algorithm has been applied in a Bayesian framework via tree-alignment space sampling.

Here we propose a new progressive algorithm to estimate an MSA under the explicit model with substitutions and indels. We have re-framed the original PIP equations into a dynamic programming (DP) approach. It aligns two MSAs (represented by their homology paths on the tree) by maximum likelihood (ML) in polynomial time. The progressive algorithm traverses a guide tree in post order; at each internal node the DP is applied to align the two sub-alignments at the child nodes. The procedure terminates at the root of the guide tree, with the complete MSA and the corresponding likelihood, which by construction is the likelihood under the PIP model. We have implemented the progressive MSA algorithm in a prototype program and verified its correctness by simulation. To our knowledge, this is the first progressive MSA algorithm using a rigorous mathematical formulation of an indel process and with polynomial time complexity.

The remainder of this manuscript is organized as follows. We first introduce notation and the PIP model. Then, we describe our DP algorithm and provide the simulation results.

## 2 The PIP model

Let 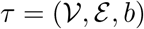 represent a rooted binary phylogenetic tree with *N* leaves. *τ* is a directed, connected, labelled acyclic graph, with a finite set of branching points 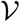 of cardinality 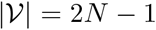 and a set of edges 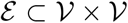. Leaves 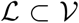 denotes *N* observed taxa, represented by strings of characters from a finite alphabet Σ (nucleotides, amino acids or codons). There are *N* − 1 internal vertices 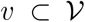 whereof the root Ω is the most recent common ancestor of all leaves. Branch length *b*(*v*) associated with node 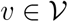 spans from *v* to its parent node pa(*v*). The total tree length ||*τ*|| is a sum of all the branch lengths.

The PIP model describes a string-valued evolutionary process along the branches of *τ*. We denote the Lebesgue measure on *τ*, i.e. the distance from the root to a given point on the tree, by the same symbol *τ*. Atomic insertions are Poisson events with rate measure *ν*(d*t*) = *λ*(*τ*(d*t*) + *μ*^−1^*δ*_Ω_(d*t*)), where *λ* is the insertion rate, *μ* the deletion rate, and *δ*_Ω_(·) Dirac’s delta function, which is one at the root Ω and zero everywhere else. This formulation guarantees that the expected sequence length remains constant during the whole evolutionary process. Point substitutions and deletions are modelled by a continuous-time Markov process on Σ_*∊*_ = Σ ⋃ *∊*, where *∊* is the deletion (or gap) symbol. Accordingly, the process generator matrix **Q**_*∊*_ of the combined substitution and indel process extends the instantaneous substitution rate matrix **Q** by a row and a column to include *∊*, which is modelled as an absorbing state as there can be no substitutions after a deletion event. The quasi-stationary distribution of **Q**_*∊*_ is denoted by *π*^*∊*^. Root Ω has a virtual infinite length stem, reflecting the equilibrium steady state distribution of the characters at the root.

The probability *ι*(*v*) of inserting a single character on branch pa(*v*) → *v* is proportional to branch length *b*(*v*). For *v* ≠ Ω it is given by *ι*(*v*) ≠ *b*(*v*)/(*τ* + *μ*^−1^); at the root atomic mass point probability *ι*(Ω) = *μ*^−1^/(||*τ*|| + *μ*^−1^) so that 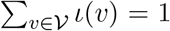. The survival probability *β*(*v*) associated with an inserted character on branch pa(*v*) → *v* is given by *β*(Ω) = 1 and *β*(*v*) = (1 *−* exp(−*μb*(*v*)))/(*μb*(*v*)).

The marginal likelihood *p_τ_* (*m*) of MSA *m* of length |*m*| is computable in *O*(*N* · |*m*|) and can be expressed as

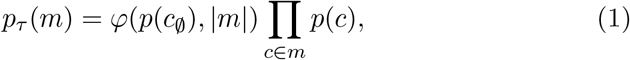

where *p*(*c*) is the likelihood of a single column *c*, and *p*(*c*_∅_) is the likelihood of an unobservable character history, represented by a column *c*_∅_ with a gap at every leaf. The factor in (1)

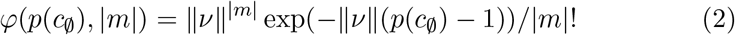

is the marginal likelihood over all unobservable character histories, where ||*ν*|| is the normalising Poisson intensity.

The column likelihood can be expressed as 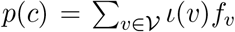, where *f_v_* denotes the probability of *c*, given that the corresponding character was inserted at *v*. This probability can be computed in *O*(*N*) using a variant of Felsenstein’s peeling recursion. Let *S* be the set of leaves that do not have a gap in column *c*, and A the set of nodes ancestral to *S*. Then

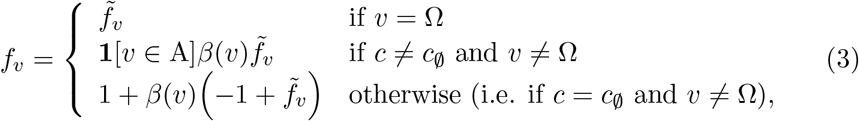

where

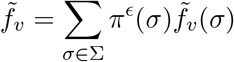

and

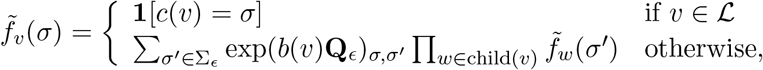

and **1**[*·*] is the indicator function.

## 3 Dynamic programming under PIP

Given an internal node *v*, our DP algorithm proceeds to align the two subalignments obtained in the left and right sub-trees maximizing the likelihood (Eq. 1) of the tree rooted at *v*. Let **X** and **Y** denote these sub-alignments, respectively with *N*_**X**_ and *N*_**Y**_ sequences and alignment lengths |**X**| and |**Y**|. If a sub-tree is a leaf then the sub-alignment, say **X**, is reduced to an input sequence, i.e. *N*_**X**_ = 1 and |**X**| corresponds the sequence length.

Note that the marginal likelihood function *p_τ_* (*m*) (Eq. 1) is not monotonically increasing in the alignment length |*m*|. While the product of column likelihoods is monotonically increasing, the marginal likelihood of unobserved histories *φ*(*p*(*c*_∅_), |*m*|) is non-monotonic. This means that *p_τ_* (*m*) cannot be maximised by means of a standard two-dimensional DP approach (in particular, because the alignment length is not known a priori). Our algorithm accounts for the dependence on alignment length with a third dimension.

The algorithm works with three three-dimensional sparse matrices **S**^M^, **S**^X^ and **S**^Y^ each of size (|**X**| + 1)×(|**Y**| + 1)*×*(|**X**| + |**Y**|) with entries defined as follows:

1. *match* cell 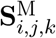 contains the likelihood of the partial optimal MSA between **X**_1_… **X**_*i*_ and **Y**_1_… **Y**_*j*_ of length *k* with the columns **X**_*i*_ and **Y**_*j*_ aligned. Consequently, all characters in the two columns are inferred to be homologous.
2. *gapX* cell 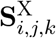 contains the likelihood of the partial optimal MSA between **X**_1_… **X**_*i*_ and **Y**_1_… **Y**_*j*_ of length *k* with the column **X**_*i*_ aligned with a column of size *N*_**Y**_ containing gaps only. The characters in the two columns do not share a common history, either because the ancestor character had been deleted on the right subtree, or because it had been inserted on the left subtree, below the node *v*.
3. similarly, *gapY* cell 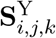 matches column **Y***j* with a column of size *N*_**X**_ containing gaps only.

### Forward phase

Each matrix **S**^M^, **S**^X^ and **S**^Y^ is initialized with *φ*(*p*(*c*_∅_)), 0) at position (0, 0, 0) and a zero in every other position. The DP equations are:

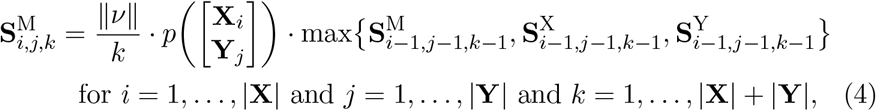

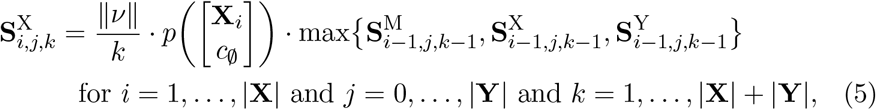

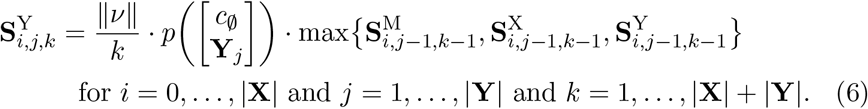

The symbol *c*_∅_ in Eq.s 5 and 6 represents a column with gaps, respectively of length *N*_**Y**_ and *N*_**X**_. The factor ||*ν*||/*k* successively constructs *φ*(*p*(*c*_∅_), *k*) along the third dimension as columns are added into partial alignments. Note that the column likelihoods *p*(·) can be computed in constant time from the corresponding column likelihoods at the two children of *v*, by re-using appropriate summands (defined by the set A in Eq. 3).

### Backtracking

An optimal alignment is determined by backtracking along a matrix **TR** of size (|**X**| + 1) × (|**Y**| + 1) × (|**X**| + |**Y**|). In the forward phase, **TR** records at position (*i, j, k*) the name of the DP matrix (“**S**^M^”, “**S**^X^”, or “**S**^Y^”) with highest likelihood at the same position (*i, j, k*). If the maximum is not unique then a uniform random choice is made. The backtracking algorithm starts at **TR**(|**X**|, |**Y**|, *k*_0_), where

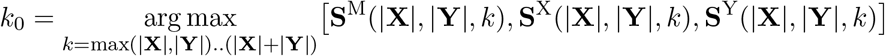

is the length of the best scoring alignment. If *k*_0_ is not unique a random uniform choice is made. **TR** is then traversed from (|**X**|, |**Y**|, *k*_0_) to (0, 0, 0). Suppose the algorithm is at position (*i, j, k*). If **TR**(*i, j, k*) = “**S**^M^” then the columns **X**_*i*_ and **Y**_*j*_ are matched and all the indices are decremented, i.e. *i* = *i* − 1, *j* = *j* − 1 and *k* = *k* − 1. If **TR**(*i, j, k*) = “**S**^X^” then the column **X**_*i*_ is matched with a column of gaps of size *N*_**Y**_ and the indices *i* and *k* are decremented, and, if **TR**(*i, j, k*) = “**S**^Y^” then the column **Y**_*j*_ is matched with a column of gaps of size *N*_**X**_ and the indices *j* and *k* are decremented.

## 4 Empirical verification of correctness

We have implemented our progressive algorithm in a prototype program. To test the correctness of algorithm and implementation, we generated data under PIP using a simulator provided by the authors 1 of PIP. We chose relatively small trees and short sequences to be able to perform analytical tests during algorithm design an program debugging. Specifically, we simulated 120 datasets in total, on trees with 4, 5, 6 and 7 leaves, using all the combinations of λ ∈ {0.1, 1} and *μ* ∈ {0.1, 1}. The resulting sequence-lengths varied between 5 and 8 nucleotides.

The simulated data was analyzed with our program using correct model parameters and guide-trees. First, we confirmed the correctness of the likelihoods obtained with the DP algorithm, by scoring the resulting MSAs with an independent implementation provided by the authors of PIP. In all cases the likelihood matched. In a second test, we verified that the DP generates optimal pairwise MSA alignments. To this end, all the possible pairwise alignments were generated at each internal node of the guide-trees and scored with the independent implementation. The DP algorithm always reconstructed an optimal MSA.

## 5 Conclusion

We have developed and implemented a progressive alignment algorithm that relies on PIP, and, thus, uses a continuous-time Markov model to describe insertions, deletions, and substitutions. The core of our method is a new DP algorithm for the alignment of two MSAs by ML, which exploits PIP’s linear time complexity (in the number of taxa and the sequence length) for the computation of marginal likelihoods. The overall complexity of the progressive algorithm is *O*(*Nl*^3^), where *N* is number of taxa and *l* the sequence length. The cubic factor stems from the fact that the likelihood is not monotonically increasing in the MSA length, so that the length has to be incorporated as an extra dimension in the DP. However, empirical findings show that the likelihood has exactly one maximum, suggesting an early stop condition to the DP. We are currently optimising our implementation in this and other time-critical aspects.

## Acknowledgement

We thank Alexandre Bouchard-Côté for providing his code to simulate sequences and to compute marginal likelihoods under PIP. This work was supported by the Swiss National Science Foundation (SNF) grant 31003A 157064 to M. Anisimova.

## Footnotes

1 personal communication

